# Prioritizing Annotated miRNAs: Only a Small Percentage are Candidates for Biological Regulation

**DOI:** 10.1101/2022.10.18.512653

**Authors:** Krystal C. Johnson, Samantha T. Johnson, Jing Liu, Yongjun Chu, David R. Corey

## Abstract

The potential for miRNAs to regulate gene expression remains controversial. DROSHA initiates the biogenesis of miRNAs while Argonaute (AGO) and TNRC6 proteins form complexes with miRNAs that recognize RNA. Here we investigate the fate of miRNAs in the absence of critical RNAi protein factors. Knockout of *DROSHA* expression reduced levels of some miRNAs, but not others. Knocking out AGO proteins, which directly contact the mature miRNA, decreased expression of miRNAs. Quantitative analysis indicates compensation to maintain the overall pool of AGO after knockout of AGO variants. Evaluation of miRNA binding to AGO proteins revealed that association between AGO and miRNAs was similar for AGO1 - 4. Contrary to the assumptions underlying many peer-reviewed reports, not all annotated miRNAs have equal potential as biological regulators. Cellular abundance, DROSHA dependence, and physical association with AGO must be considered when forming hypotheses related to their function. Our data prioritize sixty miRNAs – under two percent of the overall annotated miRNA repertoire – as being most likely to function as robust gene regulators. Our approach will facilitate identifying biologically active miRNAs.

## Introduction

MicroRNAs (miRNAs) are small, ~21 nucleotide, RNA molecules that regulate gene expression (Lee et al. 1993; Pasquinelli et al. 2000; Lee et al. 2001; Lau et al. 2001; Elbashir et al. 2001; Lagos-Quintana et al. 2001; Treiber et al. 2019). miRNAs are transcribed as longer primary miRNAs (pri-miRNAs) by RNA polymerase II (Lee et al. 2002; Lee et al. 2003; Lee et al. 2004). After transcription, an RNase III enzyme, DROSHA, initiates canonical miRNA biogenesis by forming a complex known as Microprocessor. DROSHA, in concert with its essential cofactor DGCR8, cleaves pri-miRNAs into precursor miRNAs (pre-miRNAs) in the nucleus (**Figure 1A**) (Lee et al. 2003; Gregory et al. 2004). Once the pre-miRNAs are exported to the cytoplasm, a second RNase III enzyme, DICER, along with its cofactor TRBP, processes the pre-miRNA into a mature duplex miRNA (Bernstein et al. 2001; Hutvagner et al. 2001; Rossi 2005).

**Figure 1.**
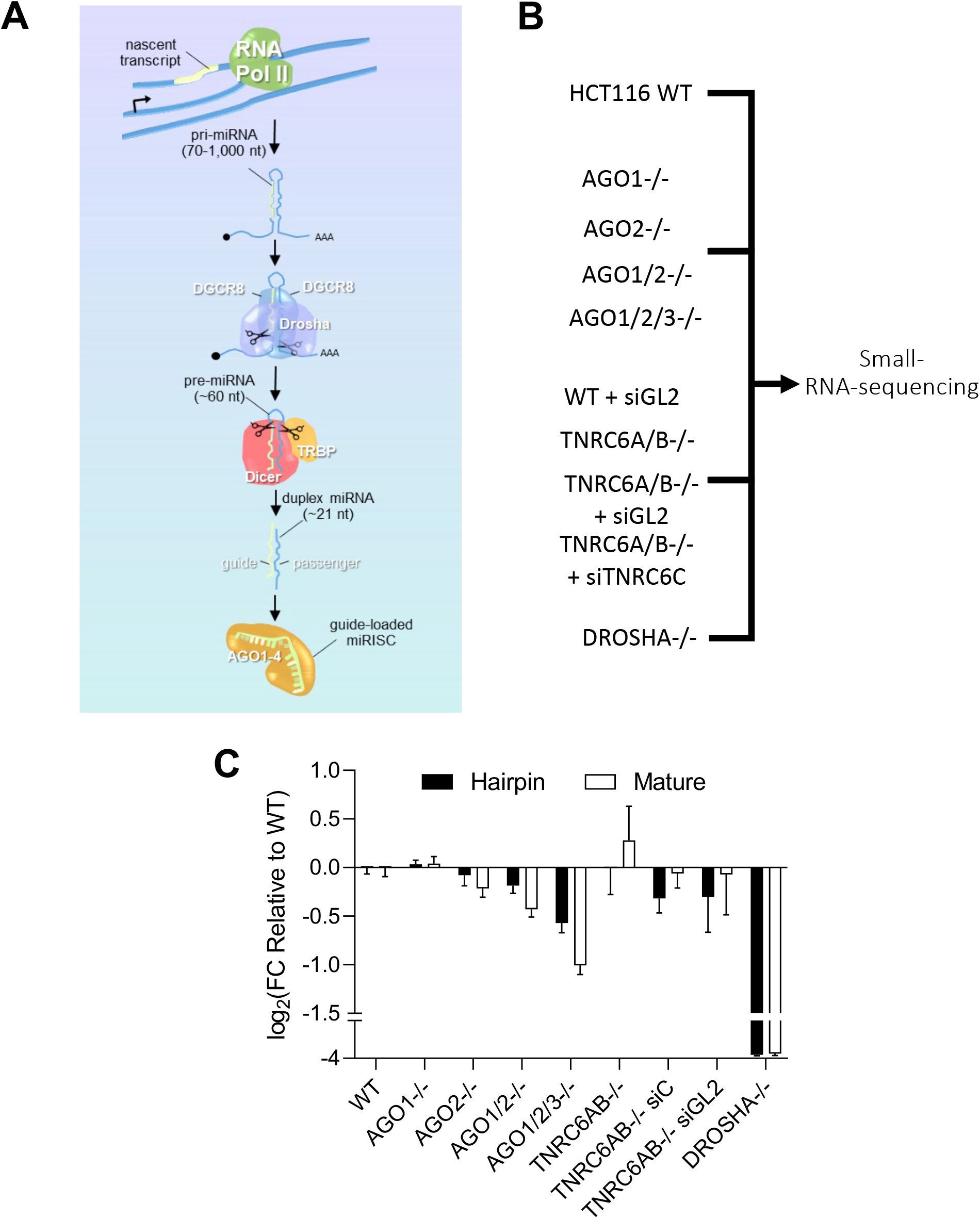
Effect of depleting essential RNAi factors on cellular miRNA profile. (A) microRNA (miRNA) biogenesis and miRNA guide strand incorporation into RNA-induced silencing complex (miRISC). (B) Experimental Scheme for analyzing HCT116 WT and knockout cell lines by small-RNA-sequencing (RNAseq) (read lengths from 17-40 nt). (C) Global hairpin and mature reads from RNAseq. Error bars represent S.D. of three biological replicates.

DICER, TRBP, PACT, and chaperone proteins load the duplex miRNA into argonaute (AGO) proteins (Chendrimada et al. 2005; Lee et al. 2006; Noland et al. 2011; Nakanishi 2016). During this loading process, the passenger strand is removed from the RNA duplex and the guide strand forms an RNA-protein complex with AGO that can efficiently and sequence-selectively bind RNA targets inside cells (MacRae et al. 2008; Yoda et al. 2010; Iwasaki et al. 2010; Chandradoss et al. 2015).

While the AGO:miRNA complex is sufficient to recognize target RNAs, recruitment of effector proteins helps modulate biological function. The miRNA:AGO complex binds to a scaffolding protein, TNRC6, through interactions between TNRC6 and AGO. TNRC6 has the potential to bind other proteins, act as a hub for protein interactions, and contribute to the control of biological functions by miRNAs (Huntzinger et al. 2013; Duchaine and Fabian et al. 2019). There are three human paralogs in human cells, TNRC6A, TNRNC6B, and TNRC6C that are related to the drosophila protein GW182. Although TNRC6 does not directly interact with the miRNA, it is needed to recruit proteins to assemble the high molecular weight RISC complex to repress the translation of the target mRNA and accelerate the mRNA’s decay (Eulalio et al. 2008; Fabian et al. 2009; Djuranovic et al. 2012; Braun et al.2013). One TNRC6 protein can interact with as many as three AGO proteins to increase dwell-time on a target mRNA and facilitate cooperativity in miRNA-mediated gene silencing (Elkayam et al. 2017; Hicks et al. 2017).

While the interactions of miRNAs and proteins associated with RNAi are well-characterized, the molecular mechanism of miRNAs in human cells is less defined. Indeed, even the number of miRNAs in the human genome and their biological relevance remains in dispute (Fromm et al. 2022). Human microRNAs were first reported over twenty years ago. Since that time the number of annotated microRNAs has increased to 2654 in miRBase (Kozomara et al. 2019).

The number of different miRNAs, when combined with the apparent simplicity and generality of seed sequence recognition, have led to many publications describing the identification of miRNAs as biomarkers and as regulators of gene expression. Understanding the mechanism of miRNA action and the application of miRNAs to therapy has not kept pace. While miRNAs have been the subject of intense study, their precise molecular mechanisms of action and the scope of their function remain undetermined (Kilikevicius et al. 2022). Lack of mechanistic understanding undermines confident interpretation of many published linkages of miRNA expression with biological function.

In this study, we seek to better understand the impact upstream biogenesis factors and downstream effector proteins have on steady-state miRNA levels. We used a suite of CRISPR/Cas9-mediated knockout cell lines (**Figure 1B**) and small-RNA-sequencing (RNAseq) to analyze the miRNA profiles in HCT116 cells. Even when two AGO proteins are knocked out, quantitative analysis reveals that expression compensates to maintain overall AGO levels. miRNAs bind similarly to AGO1-4, so even in the absence of AGO1 and AGO2 the capacity for miRNAs to form active RISC complexes with AGO protein is largely retained. These studies help define the identity of DROSHA-dependent miRNAs and the contributions of TNRC6 and AGO variants to controlling the cellular pool of miRNAs.

## Results

### Experimental design

We obtained knockout cell lines in an HCT116 colorectal cancer-derived background (**Figure 1B**). HCT116 cells were chosen because they are diploid, facilitating the CRISPR-Cas9-mediated knockout of multiple related RNAi factor genes simultaneously (i.e. *DROSHA, AGO1*, *AGO2*, *AGO3*, *TNRC6A*, and *TNRC6B*) (Liu et al. 2019; Chu et al. 2020; Chu et al. 2021). The expression of miRNAs in HCT116 cells is typical relative to other cultured cancer cell lines (Ghandi et al. 2019). We did not knock out AGO4 because, relative to the other AGO variants, it was much less expressed in HCT116 cells (Liu et al. 2019; Chu et al. 2020). We were also not able to obtain a cell line with all three TNRC6 protein paralogs (TNRC6A, TNRC6B, and TNRC6C) knocked out. A siRNA pool was used to knock down TNRC6C expression in the context of cells lacking TNRC6A and TNRC6B, referred to as TNRC6A/BKO + siTNRC6C (Liu et al. 2019).

miRNA expression was analyzed by small RNAseq and alignment with miRBase. Samples were analyzed in triplicate for each cell type to detect reads 17-40 nucleotides in length. The samples submitted for small RNAseq had RNA integrity numbers (RIN) greater than or equal to 9.4 (**Supplemental Table 1**). Multi-dimensional scaling (MDS) analysis for samples submitted for small RNAseq to detect miRNAs shows that biological replicates cluster together, demonstrating consistent results for each cell line (**Supplemental Figure 1**).

We first evaluated the effect of knocking out RNAi factors on hairpin and mature miRNA expression (**Figure 1C, Supplemental Table 1**). In *DROSHA* knockout cells, consistent with DROSHA’s role as the most upstream processing enzyme, hairpin and mature miRNA expression were reduced approximately 16-fold. miRNA expression showed little change in TNRC6A/BKO + siTNRC6C knockout/knockdown cells. Cells lacking only AGO1 or AGO2 showed little change in miRNA levels, consistent with redundancy of AGO function. Cells lacking AGO1/2 showed an intermediate reduction. AGO1/2/3 knock out cells showed the largest effect - a 2-fold reduction in mature miRNA levels.

### Effect of knocking out DROSHA on detection of miRNAs

Multiple computational and experimental approaches have led to different conclusions about the identity and annotation of miRNAs (Kozomara et al. 2019; Kim et al. 2021; Fromm et al. 2022), leading to uncertainty about the composition of bona fide miRNAs inside cells. A primary goal for this study was to prioritize annotated miRNAs for potential biological relevance.

To achieve this goal, we compared the expression of miRNAs in *DROSHA* knockout cells from three different cohorts: 1) The ~2600 miRNAs in miRBase (Kozomara et al. 2019) (**Figure 2A**); 2) The one hundred miRNAs identified as being most strongly associated with AGO2 using AGO2 eCLIP-sequencing (Chu et al. 2020) (**Figure 2BD**); and 3) RNAs identified as DROSHA-processing dependent or independent by Kim and co-workers (Kim et al. 2016; Kim et al. 2017; Kim et al. 2021) (**Figure 2CD**). These latter studies identified 311 miRNAs using multiple biochemical approaches.

**Figure 2.**
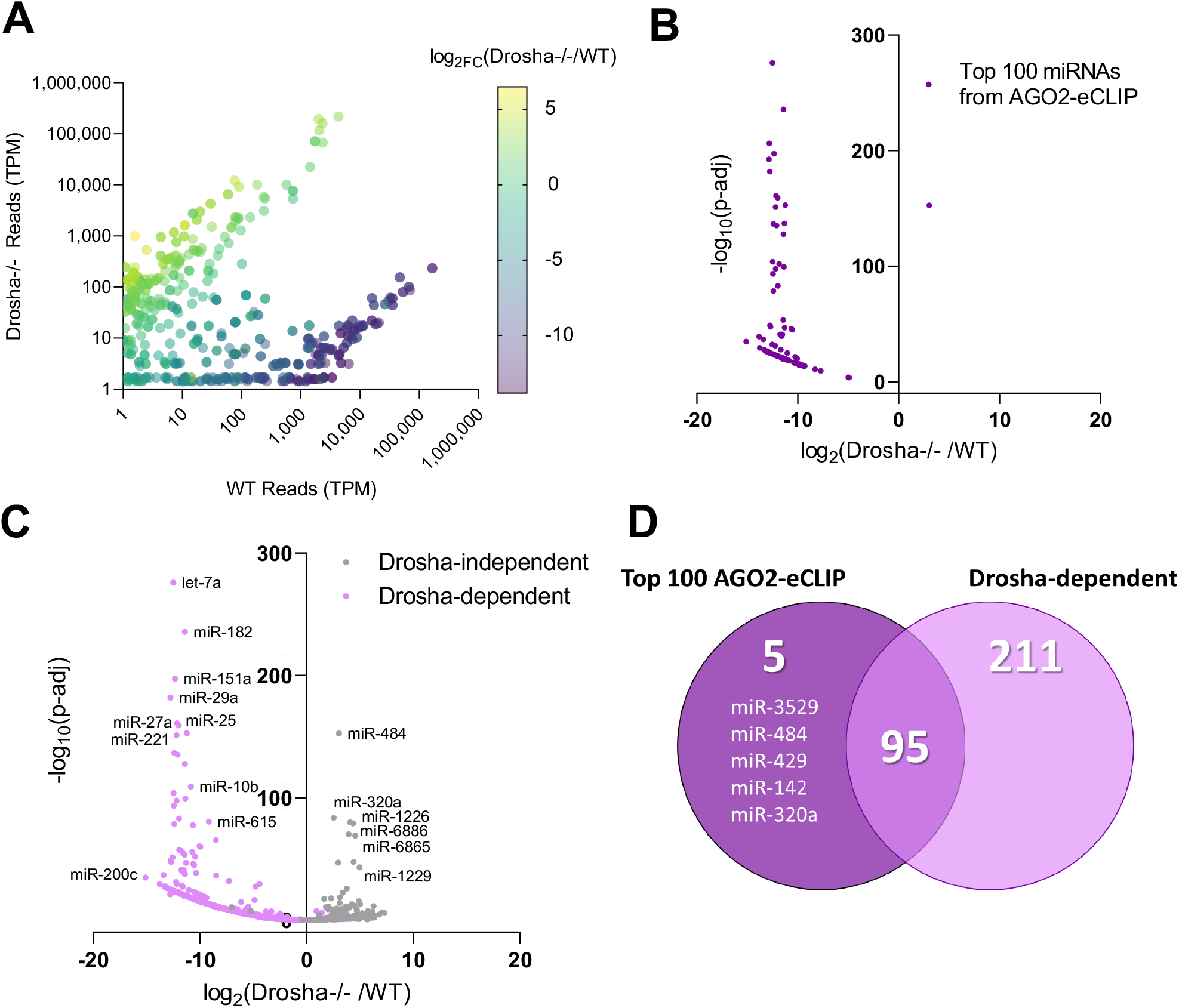
Effect of genetic deletion of *DROSHA* on mature miRNA expression. (A) Plot of global differential miRNA expression from RNAseq of *DROSHA*−/− relative to WT HCT116 cells with miRNA read cutoff at one transcript per million (N=3). (B) Differential expression in *DROSHA*−/− relative to WT cells from small-RNAseq showing only the top 100 miRNAs associated with AGO2 from AGO2-eCLIP in WT HCT116 cells. Differential expression and significance (adjusted *p*-value) calculated with DESeq2. (C) Volcano plot of differential miRNA expression in *DROSHA*−/− relative to WT where gray dots represent miRNAs that were reported as DROSHA-independent (257 total, Kim et al., 2021) and pink dots represent miRNAs that were reported as DROSHA-dependent (311 total, Kim et al., 2021). (D) Venn diagram showing overlap between miRNAs highly associated with AGO2 (Top 100 AGO2-eCLIP, Chu et al., 2020) and miRNAs that were reported DROSHA-dependent (311 pri-miRNAs reported dependent, Kim et al., 2021).

Contrary to the expectation that loss of *DROSHA* expression should abolish expression of most miRNAs, analysis of all miRNAs annotated in miRBase revealed that knocking out *DROSHA* expression led to both increased and decreased detection of annotated miRNAs (**Figure 2A**). The observation that *DROSHA* knockout increases detection of many annotated miRNAs raises the question of whether the up-regulated miRNAs were likely to be biologically relevant.

To begin to answer this question, we examined the expression levels of the miRNAs. Like any biomolecule, miRNAs that are more highly expressed will be more likely to exert a biological effect than those that are not well expressed (Cech and Steitz 2014). For reference, in wild-type cells the ten most highly detected miRNAs were characterized by 170,000-200,000 transcripts per million (**Figure 2A**, **Supplemental Figure 2A**). Of the 370 miRNAs that showed increased expression in the absence of DROSHA, most were detected at relatively low levels (less than 200 transcripts per million) in wild-type HCT116 cells (**Supplemental Figure 2B**). Only seven annotated up-regulated miRNAs were detected with transcript numbers greater than 1,000 and none were greater than 5,000 transcripts per million.

miRNAs that physically associate with AGO2 are more likely to impact biological regulation. To further evaluate the potential of miRNAs to be biologically active, we evaluated the 100 miRNAs that were most strongly associated with AGO2 protein in our eCLIP analysis (Chu et al., 2020). Almost all (98/100) miRNAs associated with AGO2 showed reduced expression in *DROSHA* knockout cells (**Figure 2B**).

We then compared our data with experimentally derived data published by Kim and co-workers on DROSHA protein-mediated processing of thousands of annotated miRNAs. Almost all miRNAs identified by Kim as DROSHA-dependent were down-regulated in *DROSHA* knockout cells (**Figure 2C**). The four DROSHA-dependent miRNAs that were upregulated in *DROSHA* knockout cells were expressed at low levels, less than 100 reads per million in wild-type cells (**Supplemental Figure 2CD**). Comparison of miRNAs from our AGO2-eCLIP and data from Kim and co-workers revealed that 95 of the top 100 AGO2-eCLIP miRNAs were also among the 311 experimentally identified DROSHA-dependent RNAs (**Figure 2D**). Taken together, these data suggest that DROSHA-dependent miRNAs in wild-type cells are also characterized by physical association with AGO2 and abundant expression.

Of the five miRNAs from the top one hundred AGO2-eCLIP list that had not previously been identified as DROSHA-dependent by Kim and coworkers, miR-3529 and miR-429 had been identified as “not yet determined” (**Supplemental Figure 2EFG**). These two miRNAs showed a statistically significant decrease in expression in *DROSHA* knockout relative to wild-type cells, suggesting that they may be DROSHA-dependent. Two other miRNAs, miR-484 and miR-320a showed a dramatic increase in expression, suggesting that they are regulated by DROSHA-independent pathways that respond to the loss of global miRNA expression.

### Effect of depleting TNRC6 paralogs on detection of miRNAs

We evaluated miRNA levels in cells lacking TNRC6 proteins. TNRC6 is a scaffolding protein responsible for bringing proteins together to regulate gene expression (Braun et al. 2013). TNRC6 is not a biogenesis factor like DROSHA, nor does it come into direct contact with miRNAs like the AGO proteins do (**Figure 3A**). It is capable of bridging two to three AGO proteins (Elkayam et al. 2017; Hicks et al. 2017), using cooperative interactions to increase the ability of the complexes between miRNAs and AGO to stably associate with target mRNAs. We evaluated miRNA expression in TNRC6A/B double knockout and triple TNRC6A/B knockout + siTNRC6C cells (**Figure 1B**).

**Figure 3.**
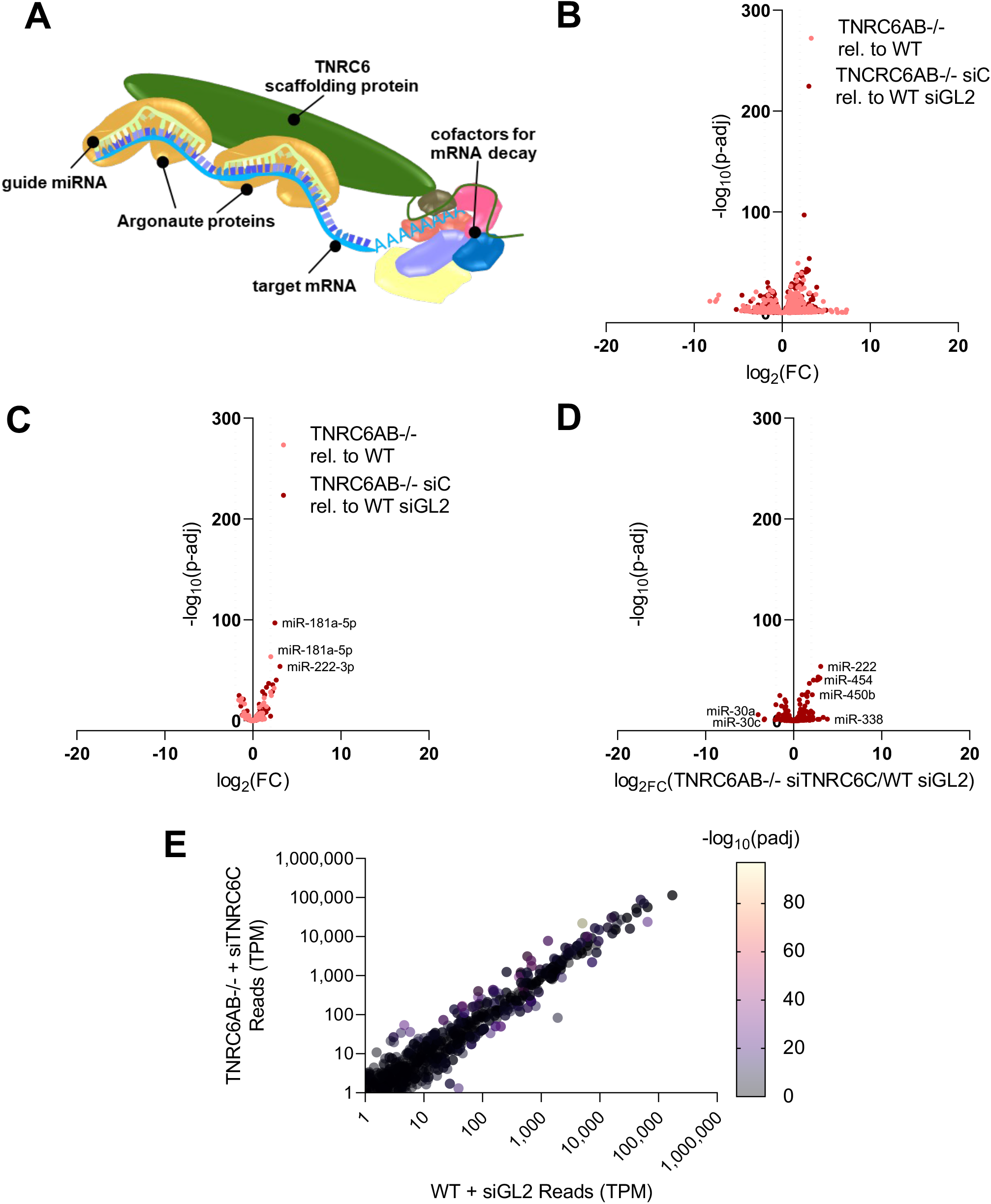
Effect of loss of TNRC6 paralogs on mature miRNA expression. (A) Scheme showing interactions of TNRC6 protein during miRNA-mediated silencing. (B) Volcano plot of global differential miRNA expression relative to WT from small-RNAseq in *TNRC6AB*−/− and *TNRC6AB*−/− siTNRC6C cells (N=3). (C) Volcano plot of differential miRNA expression relative to WT from RNAseq showing only the top 100 miRNAs associated with AGO2 from AGO2-eCLIP from HCT116. Differential expression and significance (p-adj) calculated with DESeq2. (D) Volcano plot showing the change in expression of the 311 Drosha-Dependent miRNAs (Kim et al. 2021) (E) Transcripts per million for all miRNAs with expression levels above the read cutoff of 1 transcript per million in WT transfected with siGL2 negative control and TNRC6AB−/− transfected with siTNRC6C. The color of each point represents significance, –log10 adjusted *p*-value.

Analysis of the full cohort of annotated miRNAs in miRBase revealed similar numbers of up- and down regulated RNAs for both TNRC6A/B double knockout and triple TNRC6A/B knockout +siTNRC6C cells (**Figure 3B**). The cohorts of the 100 miRNAs most enriched upon immunoprecipitation in our previous AGO2 eCLIP-sequencing (**Figure 3C**) or the 311 DROSHA-dependent miRNAs identified by Kim and coworkers (**Figure 3D**) showed a similarly mixed response to loss of the TNRC6 paralogs. The most abundantly expressed DROSHA-dependent miRNAs did not show significant expression change (**Supplemental Figure 2H**). The lack of significant expression change was maintained regardless of the expression level of the miRNA (**Figure 3E**). The mixed response and lack of statistical significance, regardless of miRNA cohort analyzed or miRNA expression level, suggests that expression of TNRC6 paralogs and their ability to act as a scaffolding protein for AGO and other cofactors does not affect miRNA levels.

### Quantitative effect of various AGO knockouts on overall pool of AGO protein

There are four AGO variants within human cells, AGO1-4 (Liu et al. 2004; Meister et al. 2004), and AGO proteins come into direct contact with miRNAs (**Figure 4A**). In HCT116 cells, AGO4 is expressed at much lower levels than AGO1-3 (Liu et al. 2019). Because AGO protein comes into direct contact with miRNAs, we examined the effect of knocking out AGO variants on miRNA levels.

**Figure 4.**
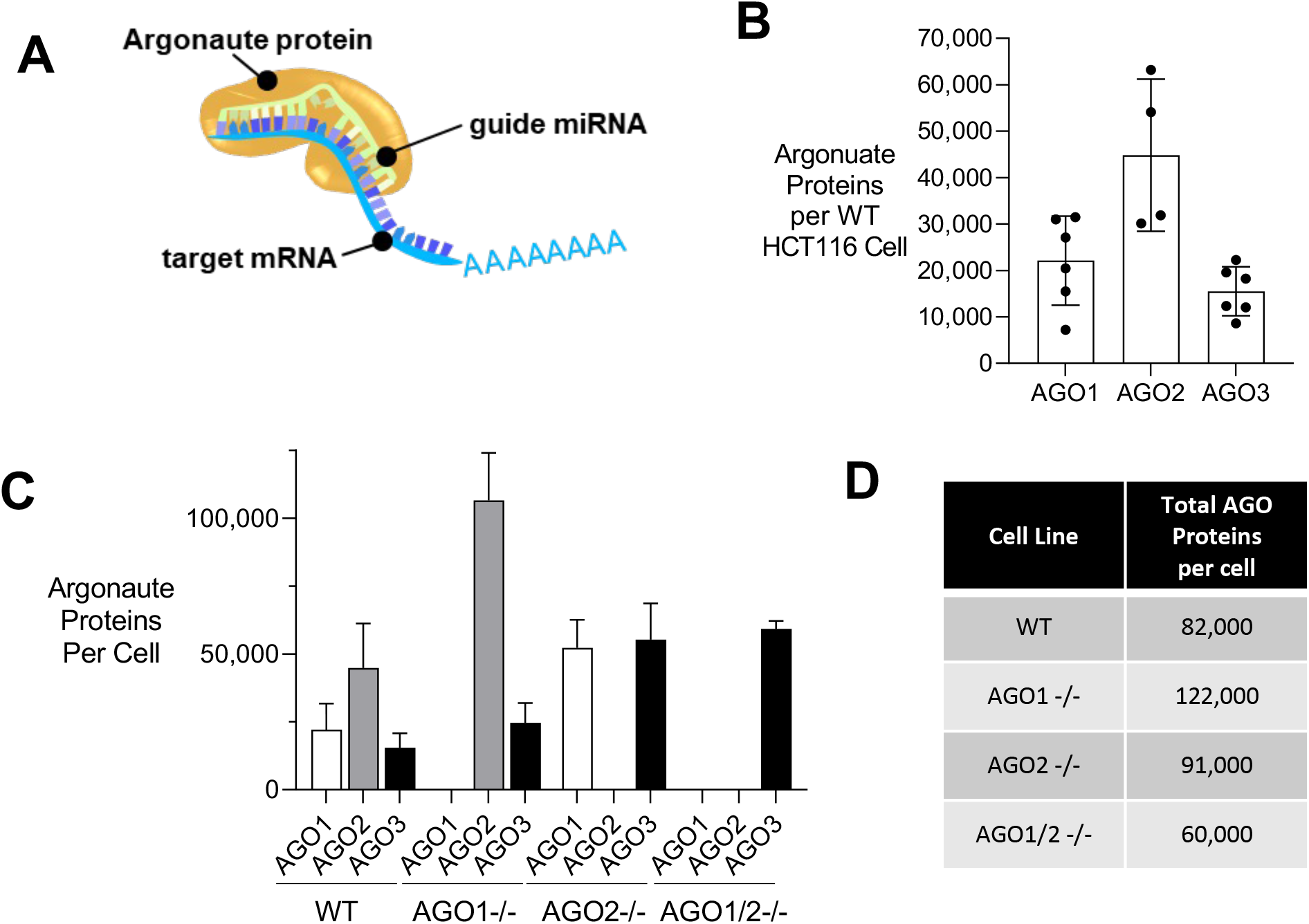
Absolute quantification of AGO proteins per cell shows compensation of remaining paralogs after AGO depletion. (A) Scheme showing the association of AGO:miRNA complex with target mRNA. (B) Absolute quantification of AGO1, AGO2 and AGO3 protein molecules in each WT HCT116 cell. (C) Absolute quantification of AGO1, 2, and 3 protein molecules per cell in HCT116 WT, *AGO1*−/−, *AGO2*−/−, and *AGO1/2*−/− cells. (D) Total AGO1, AGO2 and AGO3 protein molecules per cell in WT, *AGO1*−/−, *AGO2*−/−, and *AGO1/2*−/− HCT116 cells.

Before evaluating miRNAs, however, it was necessary to understand how many AGO protein molecules were inside cells in each of our knockout cell lines. To determine the number of AGO1-3 molecules per cell we used antibodies capable of detecting cellular AGO1, AGO2, or AGO3 proteins by western analysis in combination with titration with known quantities of recombinant AGO1, AGO2, or AGO proteins (**Supplemental Figure 3**). Using this assay, we observed 20,000, 40,000, and 15,000 copies of AGO1, AGO2, and AGO3 in each wild-type HCT116 cell (**Figure 4B**). In knockout cells, the AGO variants tended to compensate for one another. For example, when AGO2 was knocked out, AGO1 levels increased (**Figure 4C**). When AGO1 is knocked out, AGO2 levels are increased. Finally, in AGO1, AGO2, or AGO1/2 knockout cells, AGO3 levels increase. When only one or two AGO variants are knocked out, increased expression of the remaining AGO variants maintains overall AGO expression at a level near that of wild-type HCT116 cells (**Figure 4D**).

### Effect of AGO knockouts on detection of miRNAs by RNAseq

We next tested the effect of knocking out the expression of AGO proteins on miRNAs and whether that analysis would support prioritization of potentially bioactive RNAs. AGO proteins come into direct contact with miRNAs and might be expected to protect miRNAs from degradation, leading to a bigger impact on miRNA levels than the TNRC6 paralogs. We compared the effects of knocking out AGO1, AGO2, AGO1/2, and AGO1/2/3 protein expression on the expression of miRNAs.

Examining the entire catalog of annotated miRNAs in miRBase revealed that miRNA profiles were relatively unchanged when AGO1 or AGO2 were knocked out individually (**Figure 5A**). This result is consistent with previous data suggesting that AGO1 and AGO2 are at least partially redundant (Landthaler et al. 2008; Wang et al. 2012; Chu et al. 2020). There was a larger effect when AGO1/2 were knocked out together, and the largest effect was observed in the triple AGO1/2/3 knockout. By contrast to the even distribution of changes in the global miRBase miRNA catalog, we observed decreased levels for almost all one hundred most enriched miRNAs from our AGO2 eCLIP-sequencing data (**Figure 5B**). Once again, the largest effects were observed in the triple AGO1/2/3 knockout cells.

**Figure 5.**
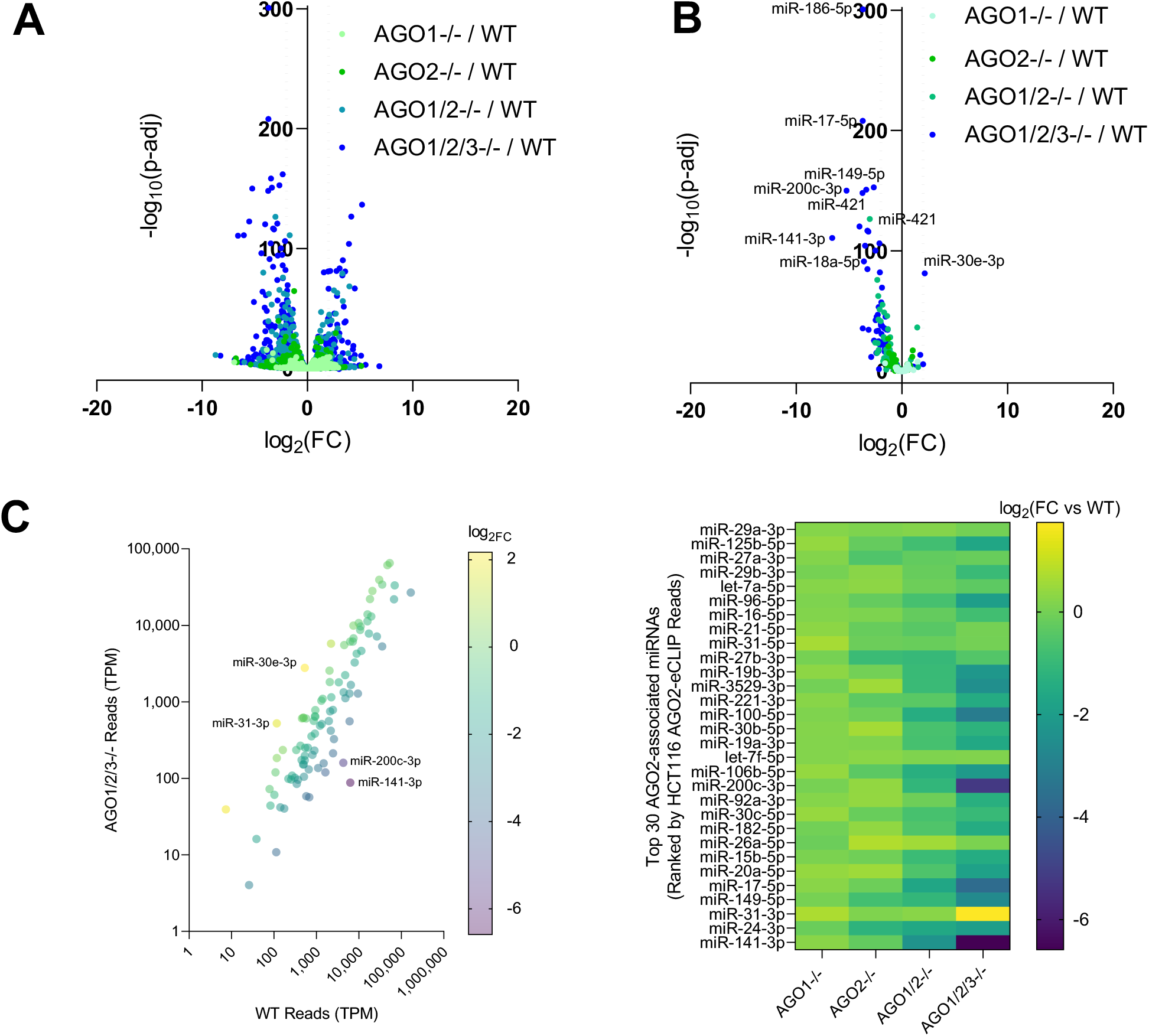
Effect of genetic deletion of AGO paralogs on mature miRNA expression. (A) Volcano plot of global differential miRNA expression relative to WT from small-RNA-sequencing of HCT116 *AGO1*−/−, *AGO2*−/−, *AGO1/2*−/− and *AGO1/2/3*−/− knockout cells. (B) Volcano plot of differential miRNA expression relative to WT from small-RNA-sequencing showing only the top 100 miRNAs associated with AGO2 from AGO2-eCLIP in WT HCT116. Differential expression and significance (p-adj) calculated with DESeq2. (C) Plot of transcripts per million for the top 100 miRNAs from AGO2-eCLIP in WT and AGO1/2/3−/− with the color of each point representing log2 fold-change. (D) Heat map showing differential expression of the top 30 miRNAs associated with AGO2 from AGO2-eCLIP in HCT116 *AGO1*−/−, *AGO2−/−*, *AGO1/2−/−* and *AGO1/2/3−/−* relative to WT.

AGO2 has been shown to be critically important for the non-canonical biogenesis of a subset of miRNAs. Two erythrocyte-specific miRNAs, miR-451 and miR-486, require AGO2 slicing for their maturation (Cifuentes et al. 2010; Cheloufi et al. 2010; Jee et al., 2018). AGO2, rather than DICER, cleaves the precursor stem of miR-451, and the mature and functional form of miR-451 is formed after additional trimming by poly(A)-specific ribonuclease (Cifuentes et al. 2010; Cheloufi et al. 2010). After DICER processing, miR-486 duplex requires AGO2 cleavage for the removal of the passenger strand (Jee et al. 2018). We found that miR-451A and miR-486 expression are low, less than 50 transcripts per million, in wild-type HCT116 cells, yet the mature form of miR-451A is decreased by 8-fold in HCT116 AGO2 knockout cells whereas miR-486 has no significant expression change. These findings support that miR-451 and miR-486 expression are cell-context specific and limited outside of the hematopoietic system, and the decrease in mature miR-451A in the absence of AGO2 reinforces the requirement for the conservation of AGO2 slicing activity in mammals.

Examination of the relative abundance of the top 100 AGO2-eCLIP miRNAs showed that the most highly expressed miRNAs in WT cells with over 1,000 transcripts per million detected were almost all downregulated or unchanged (**Figure 5C**). Evaluation of the thirty most enriched microRNAs showed the least change in the single AGO1 or AGO2 knockout cells, greater change in the double AGO1/2 knockout cells, and the greatest change in the triple AGO1/2/3 knockout cells (**Figure 5D**). Of these thirty most enriched RNAs, twenty-nine were detected at lower levels when AGO was knocked out. The largest changes observed in AGO1/2/3 knockout cells supports the absolute quantification of AGO proteins and demonstrates that the greatest loss of miRNA expression only occurs once most of the cellular pool of AGO protein is depleted.

### Association of miRNAs with AGO proteins

To further explore the impact of the association of AGO proteins and miRNAs, we used a collection of knockout cell lines and different anti-AGO antibodies (**Figures 6 and 7**) to test whether AGO variants possessed preferences for binding to miRNAs. We first immunoprecipitated cellular RNA using an anti-AGO1 or an anti-AGO2 antibody followed by small-RNAseq (**Figure 6A**).

**Figure 6.**
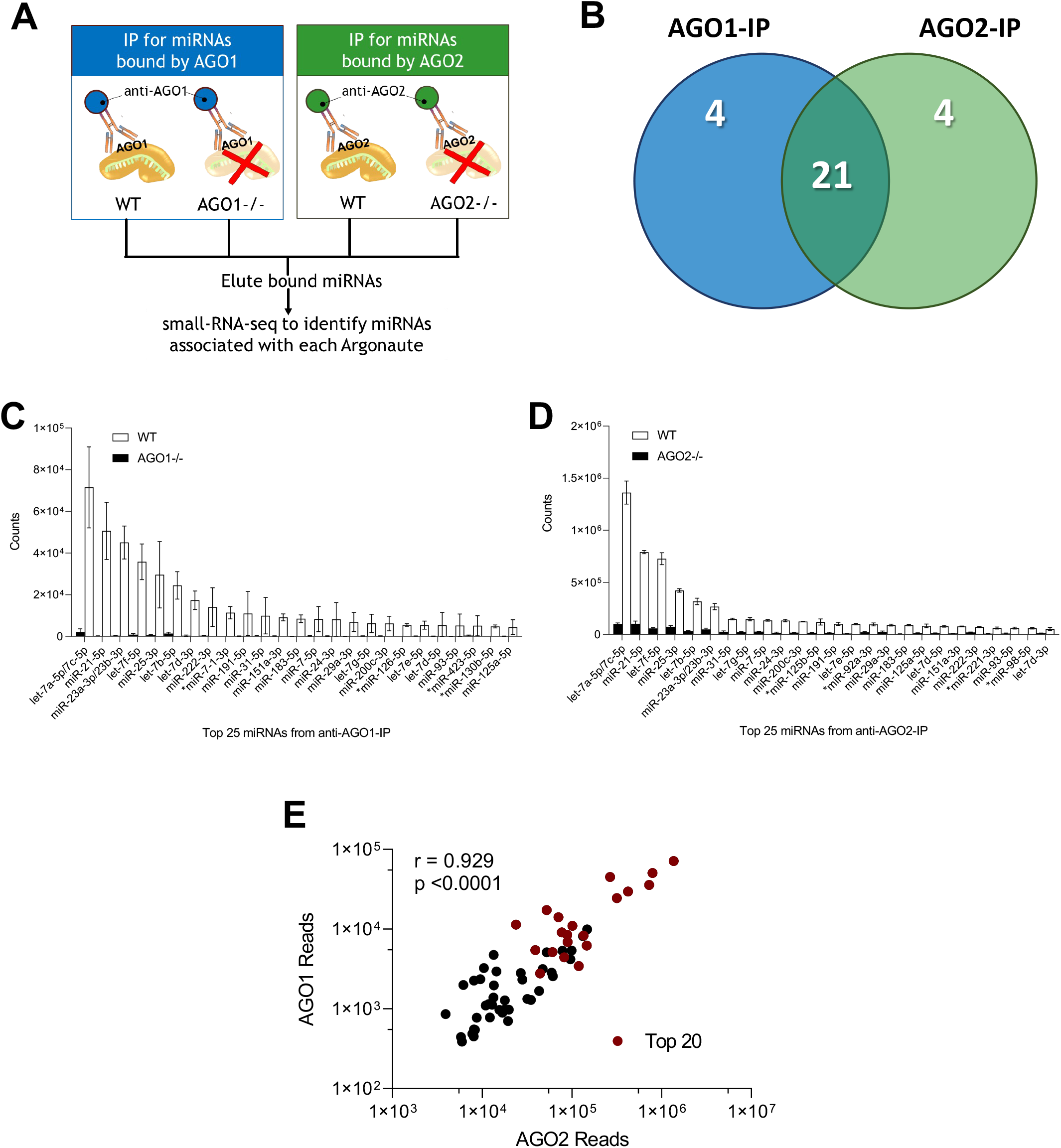
Similar abundant miRNAs found associated with AGO1 and AGO2. (A) Experimental scheme describing parallel AGO-immunoprecipitations in WT HCT116, *AGO1−/−*, and *AGO2−/−* cells. (B) Venn diagram showing the overlap between the top 25 miRNA by counts in the AGO1 and AGO2 pulldowns in the WT cell lines. (C,D) Chart showing the top 25 miRNA by counts in the (C) 2A7 Ago1 antibody pulldown in the WT and *AGO1−/−* cell lines and in the (D) 3148 anti-AGO2 antibody pulldown in the WT and *AGO2−/−* cell lines. miRNAs indicated with a (*) are the 4 miRNAs that did not overlap in both pulldowns from (B). Counts are normalized reads from DESeq2 analysis. (E) Correlation plot of miRNA reads for the top 60 miRNAs in the AGO2 and AGO1 protein pulldown in WT. The top 20 miRNAs with the highest read count from the anti-pan-AGO antibody 2A8 pulldown are highlighted in red.

**Figure 7.**
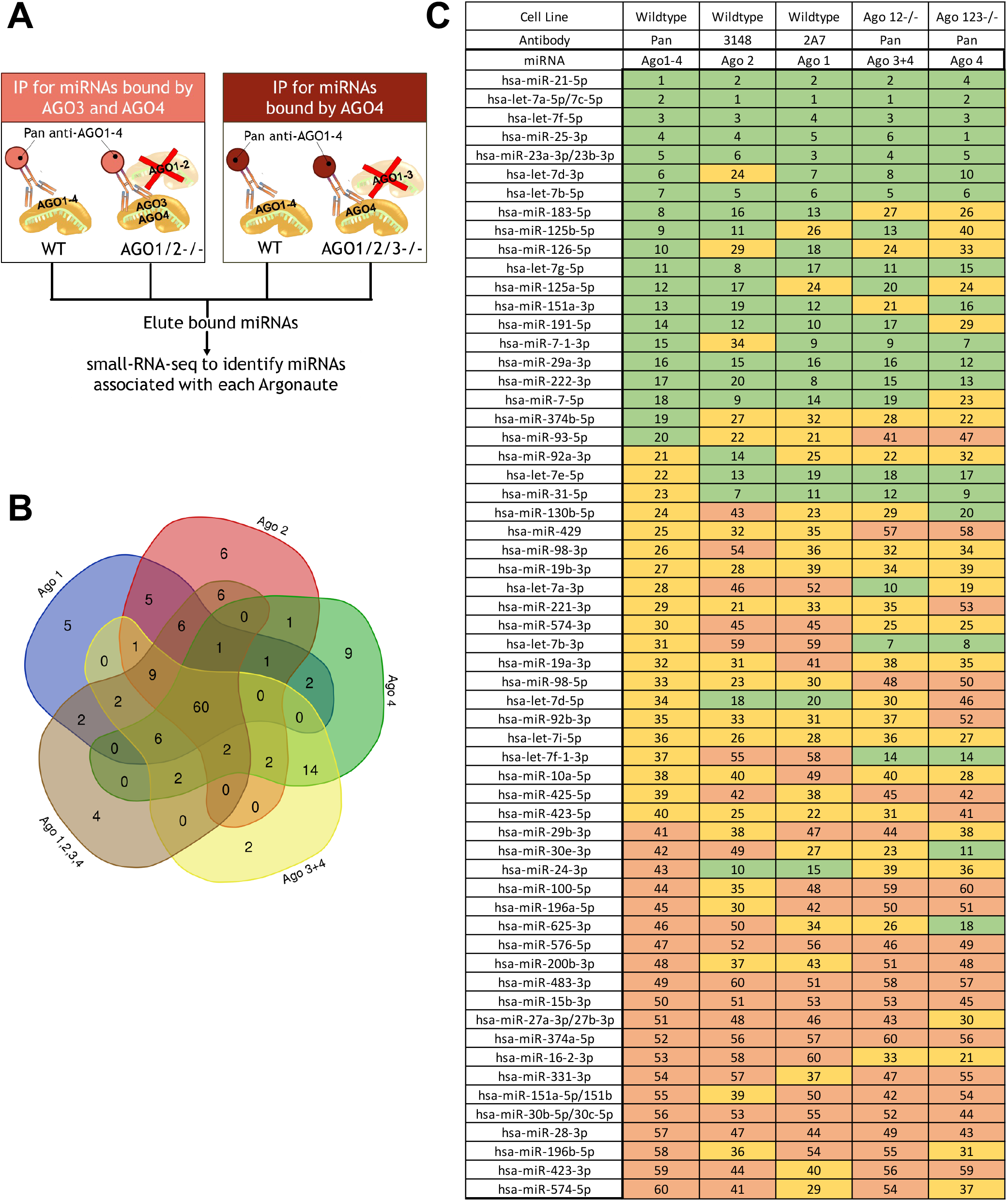
Similar abundant miRNAs found associated with all AGO proteins. (A) Experimental scheme for parallel AGO-immunoprecipitations in WT HCT116, *AGO1/2−/−*, and *AGO1/2/3−/−* cells. (B) Venn diagram of the top 100 miRNA by counts from AGO1, AGO2 and the Pan argonaute antibody pull downs in the various cell lines (WT= AGO 1,2,3,4; AGO12−/− = Ago 3,4; AGO123−/− = Ago 4). (C) Table showing where the overlapping 60 miRNAs (from Fig.7B) are ranked in each pulldown, using the Ago1-4 pulldown in the WT as a reference. The green (top 20), yellow (middle 20), and orange (bottom 20) shows the ranking of the 60 overlapping miRNAs immunoprecipitated in comparison to a previously published Ago2-eCLIP-sequencing dataset (Chu et al., 2020).

We focused on the twenty-five RNAs that were the most highly detected in each (AGO1 or AGO2) pulldown. Of the twenty-five in each cohort, twenty-one miRNAs were overlapping in both the AGO1 and the AGO2 pull-down (**Figure 6B**). The shared miRNAs were detected in a similar rank order relatively to one another (**Figure 6C and D**). A linear correlation plot for the reads from the top 60 overlapping miRNAs in the anti-AGO1 and anti-AGO2 pulldowns showed a significant Pearson correlation value (r-0.929, p<0.0001), suggesting that similar miRNAs are associated with AGO1 and AGO2 (**Figure 6E**).

While we did not possess an anti-AGO3 antibody suitable for the confident interpretation of immunoprecipitation/small RNAseq data, we did have a “PAN” AGO antibody (2A82) capable of efficiently pulling down all AGO variants (Nelson et al. 2007). The anti-AGO1, anti-AGO2, and PAN antibodies in combination with our AGO1/2 and AGO1/2/3 knockout cells lines allowed us to infer association with AGO3 and AGO4 (**Figure 6A, Figure 7A, Supplemental Figure 4A**). The immunoprecipitations performed in different AGO knockout cell lines showed nearly zero read counts compared to the same immunoprecipitation performed in WT, supporting strong enrichment of AGO-specific enrichment of associated miRNAs (**Supplemental Figure 4BC**).

Of the top one hundred miRNAs detected, 60 were detected by every anti-AGO detection scheme, indicating physical association with all four AGO proteins (**Figure 7B**). Evaluation of the IP-sequencing data based on a ranking of the reads for the associated miRNAs showed strong similarity between miRNAs in the four AGO-association cohorts (**Figure 7C**). In general, miRNAs that were in the top third of miRNAs bound to the PAN antibody in wild-type HCT116 cells were also in the top third of miRNAs in other cell types using other antibodies (**Figure 7C**). The Pearson correlation values of the different AGO-specific immunoprecipitations showed significant linear correlations (r>0.83, p<0.0001) regardless of which pair of AGO proteins were being compared demonstrating the strong overlap of miRNA identities and similar relative abundances of each miRNA associated with the different AGO proteins (**Supplemental Figure 5**). These data (**Figure 6** and **Figure 7**) suggest that AGO1, AGO2, AGO3, and AGO4 do not show strong preferences for individual miRNAs.

## Discussion

### RNA-mediated gene regulation and the necessity for prioritizing miRNAs

RNAi and small RNAs provide a powerful mechanism for controlling gene expression in mammalian cells (Gebert and MacRae 2019; Duchaine and Fabian 2019). An AGO protein binds a small RNA to form a ribonuclear protein complex in which the small RNA programs complementary recognition of target RNA sequences. The AGO protein can perform at least three critical roles: 1) protecting the RNA from degradation; 2) facilitating efficient recognition and recruitment of effector proteins for repression of target RNAs; and 3) facilitating cleavage of the fully complementary RNA sequences (in the case of AGO2 and, to a lesser extent, AGO3). The high efficiency and regulatory potential of this process is exemplified by the potent and long-lasting repression of gene expression achieved by multiple designed synthetic RNA drugs in the clinic.

While the potential of designed duplex RNAs is now well understood, the role of endogenously expressed miRNAs in cells remains less clear. While over 10,000 papers that cite the term “miRNA” are published each year (Kilikevicius et al. 2022), many of the papers that involve mammalian cells lack the exploration of mechanism necessary to confidently assign miRNAs to specific regulatory function.

There are 1917 precursor miRNAs and 2654 mature miRNAs annotated in miRbase (Kozomara et al. 2019). These thousands of potential miRNAs offer a vast scope for gene regulation. The simplicity of seed sequence pairing, which requires only seven or eight bases of perfect complementarity, fills the transcriptome with potential miRNA binding sites. Productive studies require the prioritization of miRNAs to identify those most likely to have biological impact.

### Recognition by miRNAs, not a simple process

The assumption that recognition of RNA by miRNA is a simple, predictable process drives the hypothesis behind many reports of gene regulation by small RNAs. The regulation of gene expression by miRNAs is likely to be much more complex. For canonical regulation of gene expression in the cytoplasm of mammalian cells, miRNAs bind to the 3’-untranslated region (Friedman et al. 2009). Gene regulation may involve the cooperation of multiple different miRNAs at each untranslated region, and the biological impact may be the sum of impacts from binding to several different genes. Further increasing the potential for complexity, it is also possible for miRNAs to recognize sequences within the coding region of mRNA. The need to consider the potential for multiple miRNAs to contribute to the regulation of each RNA target makes thorough prioritization of miRNAs an essential component of any plan to explore miRNA-mediated gene regulation and mechanism.

### Knockout of RNAi factors prioritize bona fide miRNAs

The goal for our study was to stratify functional miRNAs based on the impact of knocking out critical RNAi proteins on miRNA expression. DROSHA is one of the key factors responsible for processing miRNAs. We reasoned that knocking out DROSHA expression would differentiate small RNAs that were likely to be active miRNAs from small RNAs that resembled miRNAs but lacked the potential to act like miRNAs.

Of the 2654 miRNAs annotated in miRBase, 1,071 showed decreased expression when *DROSHA* expression was knocked out. Whenever genes are knocked out, however, it is always possible that secondary effects may explain altered expression when genome-wide transcription is measured. Down-regulated small RNAs might be misleadingly identified as miRNAs. Conversely, up-regulated small RNAs that are bona fide miRNAs generated by alternative biogenesis pathways might be overlooked (Yang and Lai 2011).

We used several independent approaches to build confidence in miRNA identification: 1) Since miRNA prevalence is related to biological activity, we evaluated relative expression of miRNAs and found that the more highly expressed miRNAs were also the ones most likely to be down-regulated when DROSHA was knocked out; 2) Because association with AGO is a hallmark of active miRNAs, we evaluated RNAs known to be associated with AGO2 and observed that they were also down-regulated when DROSHA was knocked out; 3) We identified miRNAs that were abundant and overlapping in association with all four AGO proteins from immunoprecipitations; and 4) Kim and coworkers performed thorough biochemical studies to identify annotated miRNAs that can be processed *in vitro* (Kim et al., 2021). Their identified DROSHA-dependent RNAs included 95/100 of the miRNAs that we had identified as most associated with AGO2. Taken together, these comparisons identify sixty mature miRNAs (**Figure 7C**) that are among the most prevalent miRNAs bound to AGO2 from AGO2-eCLIP and bound to AGO1-4 from immunoprecipitation. These data suggest these miRNAs - under two percent of the overall miRNA repertoire - as the best candidates for robust biological regulation.

We acknowledge several limitations to our approach. Identification of of “high priority” miRNAs will vary depending on cell type and environment. In some cell types, DROSHA-independent regulation is important (Cifuentes et al. 2010) and HCT116 cells are may not be an optimal model for assessing that mechanism. We focus on the most abundant miRNAs and the miRNAs that are most associated with AGO2 associated miRNAs. While these miRNAs have the most potential to act as individual gene regulators, lowly expressed RNAs may also have a significant collective impact on the control of gene networks (Ambros 2019; Chen et al. 2019).

### Link between AGO abundance and global pool of miRNAs

Absolute quantification of AGO proteins demonstrated that AGO2 is the most abundant, followed by AGO1, and then AGO3. We observed a compensatory effect for the remaining AGO proteins with enhanced AGO1, AGO2, and/or AGO3 expression to attempt to maintain the total cellular levels of AGO protein in the different AGO knockout cell lines. In support of this, we observed a modest decrease in global miRNA reads in single AGO1 and AGO2 knockout cells, a larger decrease in the AGO1/2 double knockout cells, and the greatest decrease in the AGO1/2/3 triple knockout cell line. We observed a 50% reduction in global miRNAs in the AGO1/2/3 knockout cell line. This finding supports previous studies showing that ablation of AGO2 has been linked to significant down-regulation of global miRNAs in blood and the brain (O’Carroll et al. 2007; Schaefer et al, 2010) and that loss of AGO1 and AGO2 in human melanoma cells leads to an 80% reduction in global miRNAs (Wang et al. 2012).

### Do AGO variants differ in their association with miRNAs?

We also evaluated the association of miRNAs with AGO1-4. There are four AGO variants in human cells. AGO2, and to a lesser extent AGO3, can cleave target RNA when the miRNA and target RNA form a perfect match in the enzyme active site (Wang et al. 2008; Park et al. 2017). AGO2’s slicer activity has been demonstrated to be important for endogenous siRNA biogenesis in oocytes and embryonic stem cells (Babiarz et al. 2008; Tam et al., 2008; Watanabe et al. 2008), miR-451 biogenesis in blood (Cifuentes et al. 2010), rare mRNA cleavage by miRNAs (Karginov et al. 2010), and precursor cleavage in U20S and 293 cells (Diederichs and Haber 2007). That ability to cleave RNA with higher efficiency sets AGO2 apart from the other AGO proteins, but variation in binding to miRNAs among the four paralogs was less well known.

When we compared the profile of miRNAs associated with each of the endogenous AGO proteins, we observed consistent overlap in the miRNA species after immunoprecipitation with anti-AGO antibodies (**Figure 6C**). These results support previous findings that pulled down (1) tagged AGO1-4 in HEK293 cells (Hafner et al. 2010), (2) endogenous AGO1-3 proteins in AGO1, AGO2, and AGO1/2 knockout human melanoma cells (Wang et al. 2012), and (3) endogenous AGO1-4 proteins in HeLaS3 cells (Dueck et al. 2012) and showed that overall miRNAs distribute randomly among the individual AGO proteins according to their relative protein abundance.

### Conclusions

We have evaluated the impact of *DROSHA*, *TNRC6*, and *AGO* expression on levels of miRNAs. Knockout of the key biogenesis factor *DROSHA* helps prioritize miRNAs that have potential to be involved in biological regulation. Knockout of the TNRC6 paralogs had little effect on miRNA levels, consistent with the role of TNRC6 as a scaffolding protein that organizes regulatory proteins but does not come into direct contact with miRNAs. Quantitative analysis of protein expression reveals that knocking out AGO1 and AGO2 leads to compensatory up-regulation of AGO3. Knocking out AGO1, AGO2, and AGO3 reduces the cellular pool of miRNAs with the greatest effect observed for the miRNAs physically associated with AGO proteins. The sixty proteins identified in our analysis as the most likely candidates for biological regulation bind all four AGO proteins, suggesting that regulation may be maintained even in the absence of AGO1 and AGO2. Our data explain why some miRNAs may be more likely than others to play measurable roles in regulating biological function than others and how core RNAi protein factors affect levels of active miRNAs.

## Materials and Methods

### Cell Culture

Wild-type HCT116 cells were obtained from ATCC. HCT116 cells containing knock out modifications to the DROSHA, AGO1, AGO2, AGO1/2, AGO1/2/3, TNRC6A, TNRC6B, and TNRC6A & TNRCB genes were purchased from GenScript. All cell lines were cultured in McCoy’s 5A medium (Sigma-Aldrich) supplemented with 10% FBS (Sigma-Aldrich) in 37°C 5% CO_2_.

### Transfections

All transfections used Lipofectamine RNAi MAX (Invitrogen). For transfections, cells were seeded into six-well plates at 150,000 cells per well for wild type, TNRC6A−/−, and TNRC6B−/−. TNRC6AB−/− cells were seeded at 250 thousand cells per well due to the slowed growth rate of these cells. Cells were transfected with siGL2 as a negative control and a siRNA pool targeting TNRC6C as described in Liu et al.2019.

### RNA extraction and small-RNA-seq

Whole cell RNA was extracted from cells with Trizol at 80% confluency. This whole cell RNA was submitted to the UTSW Genomics Sequencing Core for small-RNA-sequencing (also referred to as microRNA-sequencing). The miRNA libraries were prepared using Illumina TruSeq Small RNA library preparation kits (Catalog # RS-200-0012, RS-200-0024). Protocol from TruSeq® Small RNA Library Prep Reference Guide (Document # 15004197 v02, July 2016) was followed. The total RNA quality was checked with Bioanalyzer (Agilent, RNA 6000 Nano kit, 5067-1511). Small RNA library preparation started with 1ug high quality total RNA. First, the small RNAs were ligated with 3’ Adapter and then ligated with 5’ Adapter. Reverse transcription followed by amplification creates cDNA constructs based on the small RNA ligated with 3’ and 5’ adapters. Then libraries were cleaned with Agencourt AMPure beads (Beckman Coulter, Catalog # A63882). Then the concentration of the remaining clean RNA was measured by Picogreen (Fisher scientific, Quant-iT PicoGreen dsDNA Assay Kit, P7589). Equal amounts were pooled from each library. Then the library pool was further purified using Pippin Prep system (Sage Science INC) with 3% agarose gel cassettes (Sage Science INC, CDP3010). The final library pool quality was checked with Bioanalyzer (Agilent, High Sensitivity DNA Kit, 5067-4626) and qPCR (Kapa library quantification kit, 7960336001). Pooled libraries were sequenced on NextSeq v2.5 High Output flow cells as single end 50 cycle runs, the yield per library was between 6.9-16.3 million pass filter reads, with a mean quality score of 34.6 and a % >= Q30 of 95.8% bases. Cutadapt trimmed read lengths were between 25 to 40 bases, mapping reference was mature and hairpin miRNA sequences from GRCh37.

### Argonaute protein associated miRNA immunoprecipitation

HCT116 WT and AGO KO cells were grown in 150 mm2 dishes in McCoy medium with 10% FBS. Cytoplasmic and nuclear fraction were isolated and lysed by Cyto lysis buffer [20 mM Tris–HCl (pH 7.4), 150 mM NaCl, 2 mM MgCl2 and 0.5% NP-40, 0.5 mM DTT] with proteinase inhibitor and RNase inhibitor and nuclear lysis buffer [150 mM KCl, 20 mM Tris–HCl 7.4, 3 mM MgCl2, 0.5% NP-40, Roche protease inhibitors cocktail, RNAse in (50 U/ml final)]. Protein A/G agarose beads were briefly washed with corresponding lysis buffer before use. 100 mg of cytoplasmic protein or nucleus protein from WT and AGOs KO cell lines were precleared by binding with 100 μL blank protein A/G agarose beads in 500ul volume for 1 hour at 4°C. One hundred microliters of protein A/G agarose beads were incubated with 5 mg of anti-AGO1(WAKO Chemical, #015-22411), anti-AGO2 antibody (3148, gift from Jay A. Nelson lab) or pan antibody (Sigma, MABE56) in 0.5 mL at 4°C with gentle agitation for 2 h. After spin, the precleared cytoplasmic or nuclear protein supernatant was transferred into antibody binding tube and rotated ON at 4°C. After washing with lysis buffer (three times), the beads were then treated with elution buffer (1% SDS, 0.1M NaHCO_3_ and RNase inhibitor). Following proteinase K treatment, RNA was extracted by phenol:chloroform, precipitated by ethanol. RNA pellet was dissolved by nuclease free water and sent for miRNA sequencing.

### Absolute argonaute protein quantification

Recombinant AGO (AGO1, AGO2, AGO3) protein was purchased from Active Motif (AGO1 Catalog No: 31522, AGP2 Catalog No: 31886, AGO3 Catalog No: 31523). The protein concentration of each recombinant Argonaute stock was confirmed by BCA Assay (ThermoScientific, Catalog No:23225). Recombinant Argonaute protein was serially diluted and used to construct a standard curve for Western blot analysis. Serial dilutions were performed in Protein LoBind tubes (Eppendorf) coated with Bovine Serum Albumin Standard protein (ThermoScientific, 2mg/mL) to prevent protein loss. Western blot images were exposed on film and analyzed with ImageJ to construct a standard curve plotting Western signal vs recombinant protein concentration. The slope of this line was used to determine the Argonaute proteins per cell in whole cell lysate from WT, AGO1−/, AGO2−/−, and AGO1/2 −/− HCT116 cells. Whole cell lysis buffer contains 50 mM Tris pH 7.0, 120 mM NaCl, 0.5% NP-40, 1 mM EDTA, 1 mM DTT.

### Statement about statistical tests

Differential analysis of small-RNA-seq expression changes were performed using DESeq2 package to generate fold-change and adjusted p-values. Pearson correlation values and two-tailed p-values were calculated for correlations on GraphPad Prism 9.1.2.

## Data availability

The data discussed in this publication have been deposited in NCBI’s Gene Expression Omnibus (Edgar et al., 2002) and are accessible through GEO Series accession number GSE214157 (https://www.ncbi.nlm.nih.gov/geo/query/acc.cgi?acc=GSE214157) and GSE214235 (https://www.ncbi.nlm.nih.gov/geo/query/acc.cgi?acc=GSE214235). Small-RNA-seq of miRNAs from Argonaute1-4 immunoprecipitation can be accessed at GSE214157, and small-RNA-seq of whole cell miRNAs detected in the different RNAi factor knockout cell lines can be accessed at GSE214235.

## Acknowledgements

This study was supported by 1F31GM137591 (K.C.J.) and R35GM118103 (D.R.C.) from the National Institutes of Health and the Robert A. Welch Foundation I-1244 (D.R.C.). The authors thank the UTSW Genomics Sequencing Core for microRNA-seq library preparation, sequencing, and data analysis.

## Author contributions

K.C.J, S.T.J., and J.L. performed and analyzed experiments. K.C.J. and D.R.C prepared the manuscript. D.R.C. supervised the study.

---

**Supplemental Table 1.** Quality of Input RNA and Depth of Small-RNA-Sequencing. (A) Table describing mapped and unmapped hairpin/mature reads from small-RNA-sequencing of HCT116 WT, WT + siTNRC6C, WT + siGL, AGO1−/−, AGO2−/−, AGO1/2−/−, AGO1/2/3−/−, TNRC6AB−/−, TNRC6AB−/− + siTNRC6C, and Drosha−/− generated with MultiQC (N=3, each replicate shown). RNA Integrity Number for each sample is listed from Bioanalyzer measurement.

| Cell Line | RNA Integrity Number (RIN) | Hairpin Reads |  |  | Mature Reads |  |  |
| --- | --- | --- | --- | --- | --- | --- | --- |
|  |  | Mapped | Unmapped | Average Mapped | Mapped | Unmapped | Average Mapped |
| WT-1 | 10 | 1,241,453 | 2,596,056 | 1,188,383 | 2,914,161 | 3,698,136 | 2,770,099 |
| WT-2 | 9.9 | 1,190,020 | 2,063,472 |  | 2,818,701 | 3,121,801 |  |
| WT-3 | 10 | 1,133,676 | 2,778,649 |  | 2,577,435 | 3,772,466 |  |
| WT-siTNRC6C-1 | 9.7 | 924,896 | 5,002,182 | 877,331 | 2,328,480 | 5,829,641 | 2,278,721 |
| WT-siTNRC6C-2 | 9.8 | 839,177 | 4,354,293 |  | 2,200,438 | 5,102,894 |  |
| WT-siTNRC6C-3 | 9.9 | 867,920 | 5,509,666 |  | 2,307,246 | 6,285,098 |  |
| WT-siGL-1 | 9.7 | 961,881 | 3,102,320 | 1,015,762 | 2,389,127 | 3,948,630 | 2,560,588 |
| WT-siGL-2 | 10 | 1,049,884 | 3,254,220 |  | 2,715,448 | 4,182,055 |  |
| WT-siGL-3 | 9.4 | 1,035,522 | 3,432,661 |  | 2,577,188 | 4,346,786 |  |
| AGO1KO-1 | 10 | 1,246,089 | 2,927,324 | 1,216,268 | 2,908,645 | 4,025,536 | 2,853,976 |
| AGO1KO-2 | 10 | 1,176,193 | 2,124,019 |  | 2,696,887 | 3,160,239 |  |
| AGO1KO-3 | 10 | 1,226,521 | 2,283,438 |  | 2,956,396 | 3,363,881 |  |
| AGO2KO-1 | 10 | 1,229,026 | 2,175,239 | 1,128,661 | 2,559,208 | 3,291,328 | 2,389,085 |
| AGO2KO-2 | 10 | 1,091,166 | 2,165,509 |  | 2,331,469 | 3,154,923 |  |
| AGO2KO-3 | 10 | 1,065,790 | 1,901,273 |  | 2,276,577 | 2,867,319 |  |
| AGO1/2KO-1 | 10 | 1,071,086 | 2,724,271 | 1,046,534 | 2,178,486 | 3,703,268 | 2,060,852 |
| AGO1/2KO-2 | 10 | 981,581 | 3,392,742 |  | 1,945,672 | 4,287,033 |  |
| AGO1/2KO-3 | 10 | 1,086,934 | 2,327,381 |  | 2,058,399 | 3,322,827 |  |
| AGO1/2/3KO-1 | 10 | 789,422 | 3,183,939 | 801,701 | 1,324,122 | 3,905,127 | 1,382,294 |
| AGO1/2/3KO-2 | 10 | 753,284 | 2,912,430 |  | 1,332,435 | 3,597,338 |  |
| AGO1/2/3KO-3 | 10 | 862,398 | 2,783,925 |  | 1,490,326 | 3,571,722 |  |
| TNRC6ABKO-1 | 9.9 | 1,053,427 | 4,065,333 | 1,198,046 | 3,027,992 | 5,006,402 | 3,434,959 |
| TNRC6ABKO-2 | 9.4 | 1,069,406 | 5,950,065 |  | 2,832,625 | 6,899,871 |  |
| TNRC6ABKO-3 | 9.9 | 1,471,305 | 2,532,066 |  | 4,444,261 | 3,838,375 |  |
| TNRC6ABKO-siTNRC6C-1 | 9.9 | 1,065,536 | 6,098,167 | 957,097 | 2,977,508 | 7,026,554 | 2,666,311 |
| TNRC6ABKO-siTNRC6C-2 | 9.4 | 935,352 | 9,676,080 |  | 2,588,547 | 10,493,056 |  |
| TNRC6ABKO-siTNRC6C-3 | 9.9 | 870,403 | 10,262,574 |  | 2,432,878 | 11,032,480 |  |
| TNRC6ABKO-siGL-1 | 9.8 | 1,283,843 | 3,801,590 | 984,056 | 3,666,018 | 4,934,676 | 2,712,357 |
| TNRC6ABKO-siGL-2 | 9.8 | 853,069 | 3,392,062 |  | 2,310,178 | 4,144,931 |  |
| TNRC6ABKO-siGL-3 | 10 | 815,255 | 4,336,094 |  | 2,160,874 | 5,061,393 |  |
| DROSHAKO-1 | 9.6 | 83,227 | 4,074,953 | 87,629 | 197,912 | 4,148,445 | 215,102 |
| DROSHAKO-2 | 9.4 | 91,455 | 5,855,500 |  | 233,584 | 5,935,849 |  |
| DROSHAKO-3 | 9.5 | 88,206 | 4,325,593 |  | 213,811 | 4,402,731 |  |

**Supplemental Figure 1.**
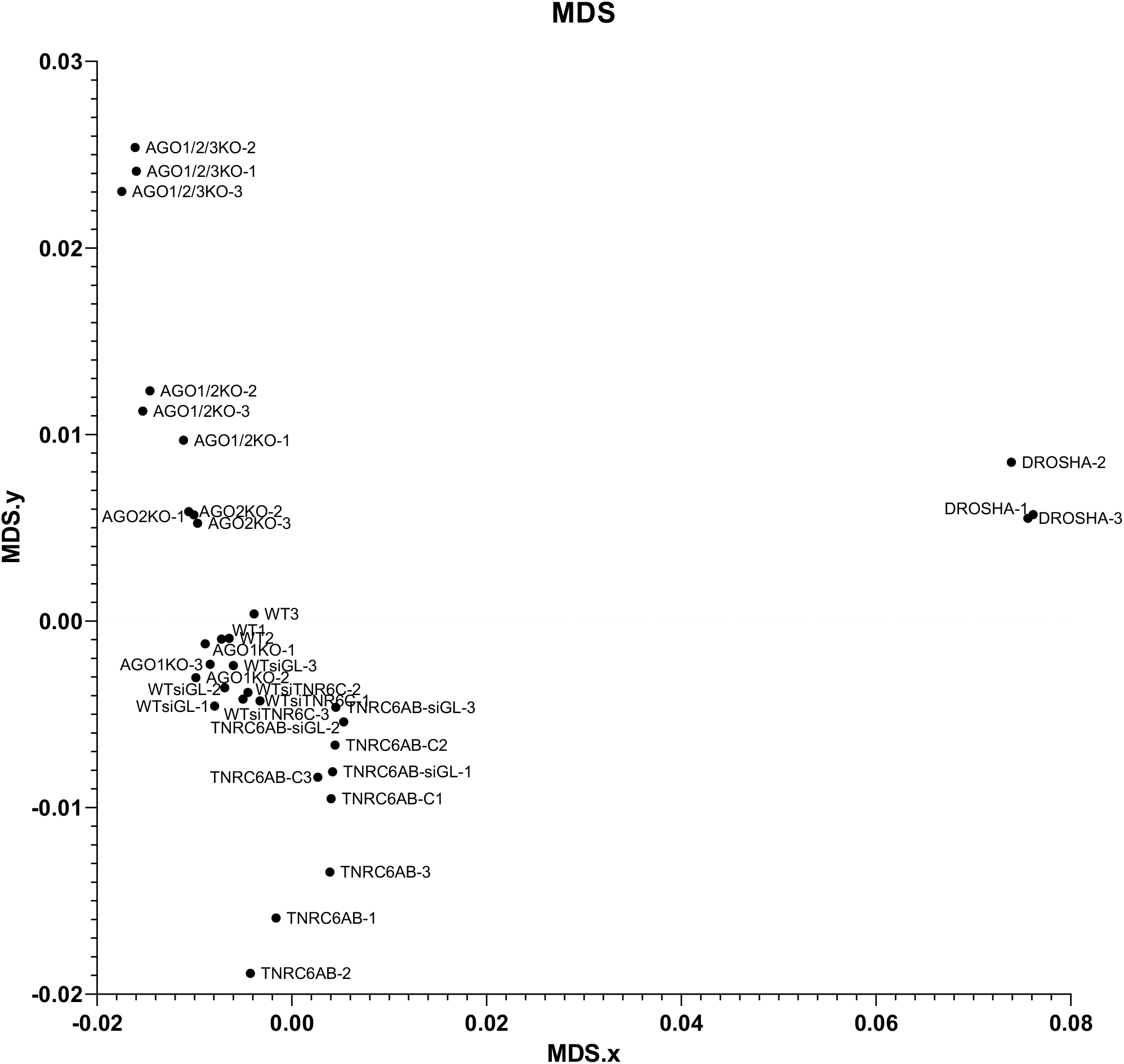
Clustering of Samples Based on Small-RNA-Sequencing Results. Plot of Multi-Dimensional Scaling (MDS) analysis describing small-RNA-sequencing of HCT116 WT, WT + siTNRC6C, WT + siGL, AGO1−/−, AGO2−/−, AGO1/2−/−, AGO1/2/3−/−, TNRC6AB−/−, TNRC6AB−/− + siGL TNRC6AB−/− + siTNRC6C, and DROSHA−/− generated with MultiQC (N=3, each replicate shown).

**Supplemental Figure 2.**
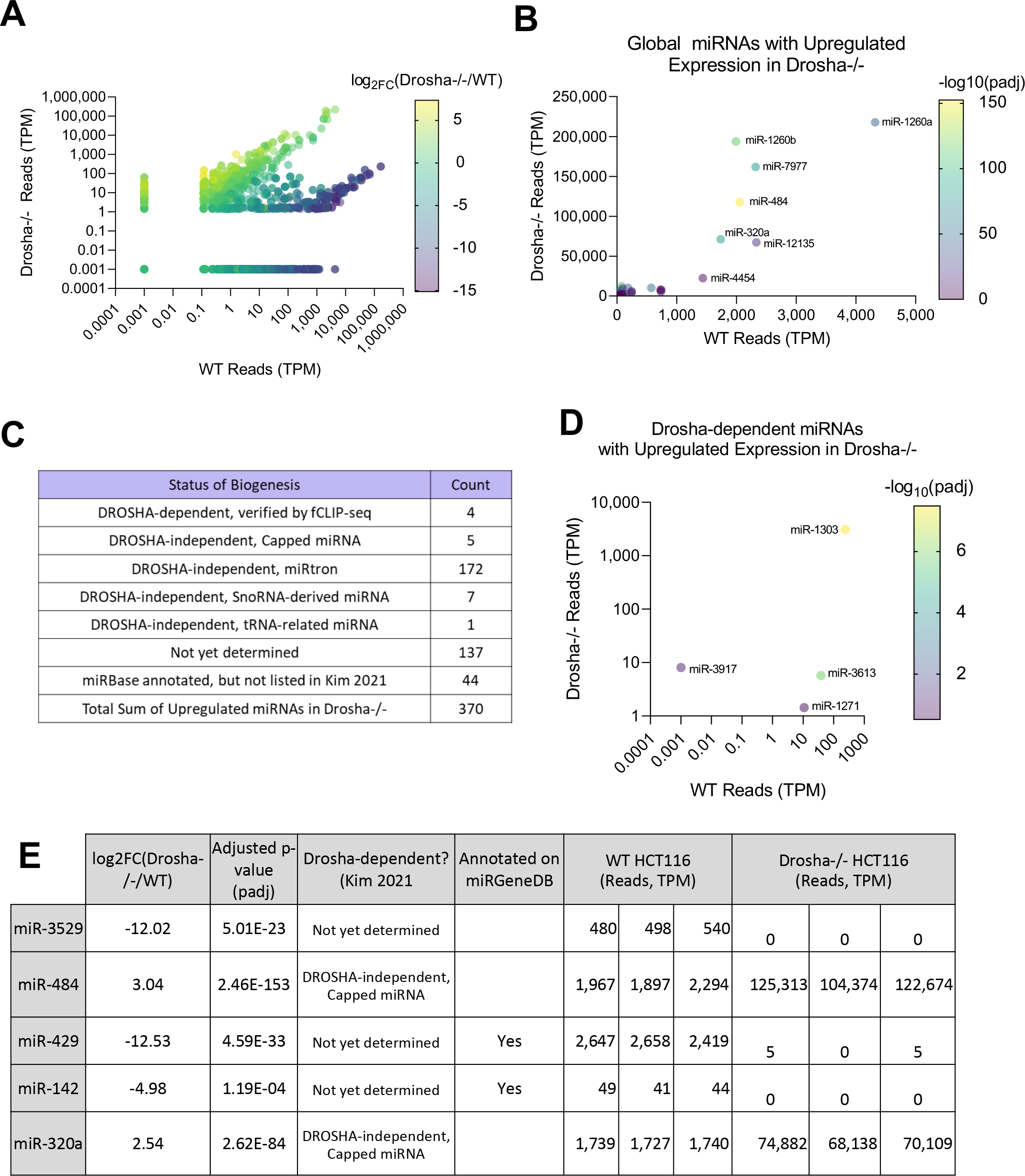
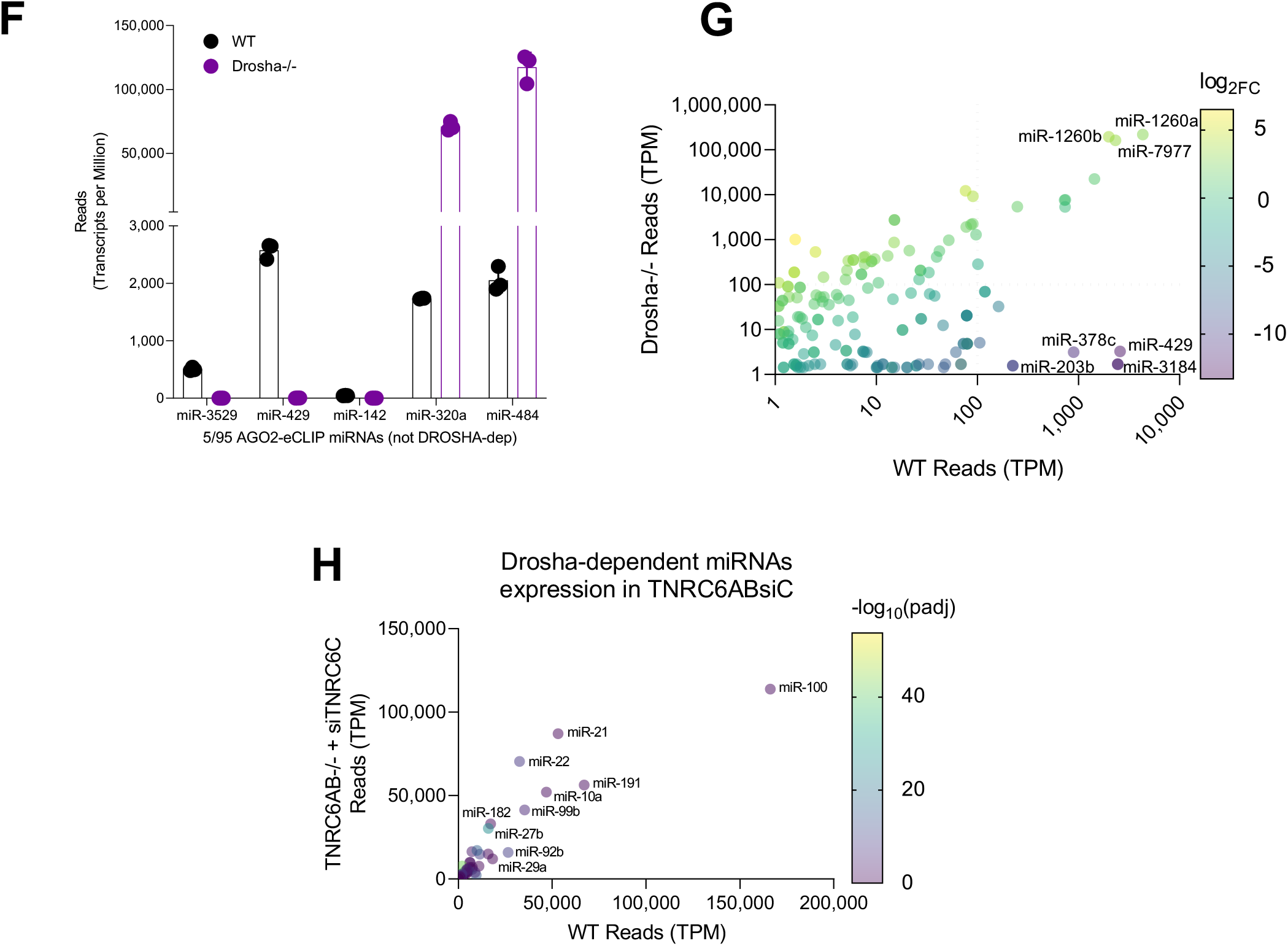
miRNA Expression in Reads in different RNAi KO Cell Lines. (A) Plot of all detected miRNA reads in transcripts per million (TPM) in DROSHA−/− and WT cells. The color of each dot represents the log2 fold-change of each miRNA in DROSHA−/− relative to WT. (B) Plot of reads for all miRNAs that were upregulated in DROSHA−/− relative to WT. (C) Reported status of miRNA biogenesis (Kim et al., 2021) for all miRNAs that were upregulated in DROSHA−/− relative to WT. (D) Plot of reads for the four DROSHA-dependent miRNAs (Supp. Fig 1C, Kim et al., 2021) with upregulated expression in DROSHA−/− relative to WT. (E) Table showing information about the 5 miRNAs highly associated with AGO2 (Top 100 AGO2-eCLIP, Chu et al., 2020) that were reported Drosha-independent or not yet determined (Kim et al., 2021). (F) Bar graph showing reads (TPM) detected for the 5 miRNAs from Table S2E in WT HCT116 and DROSHA−/− cells. (G) Plot of relative abundance and differential expression of miRNAs that were reported as not yet determined as DROSHA-dependenct (1,313 total, Kim et al., 2021). (H) Plot of detected miRNA reads for all DROSHA-dependent miRNAs (Kim et al., 2021) in TNRCAB−/− + siTNRC6C relative to WT. The color of each dot represents the significance of the miRNA expression change as negative log10 of the adjusted p-value.

**Supplemental Figure 3.**
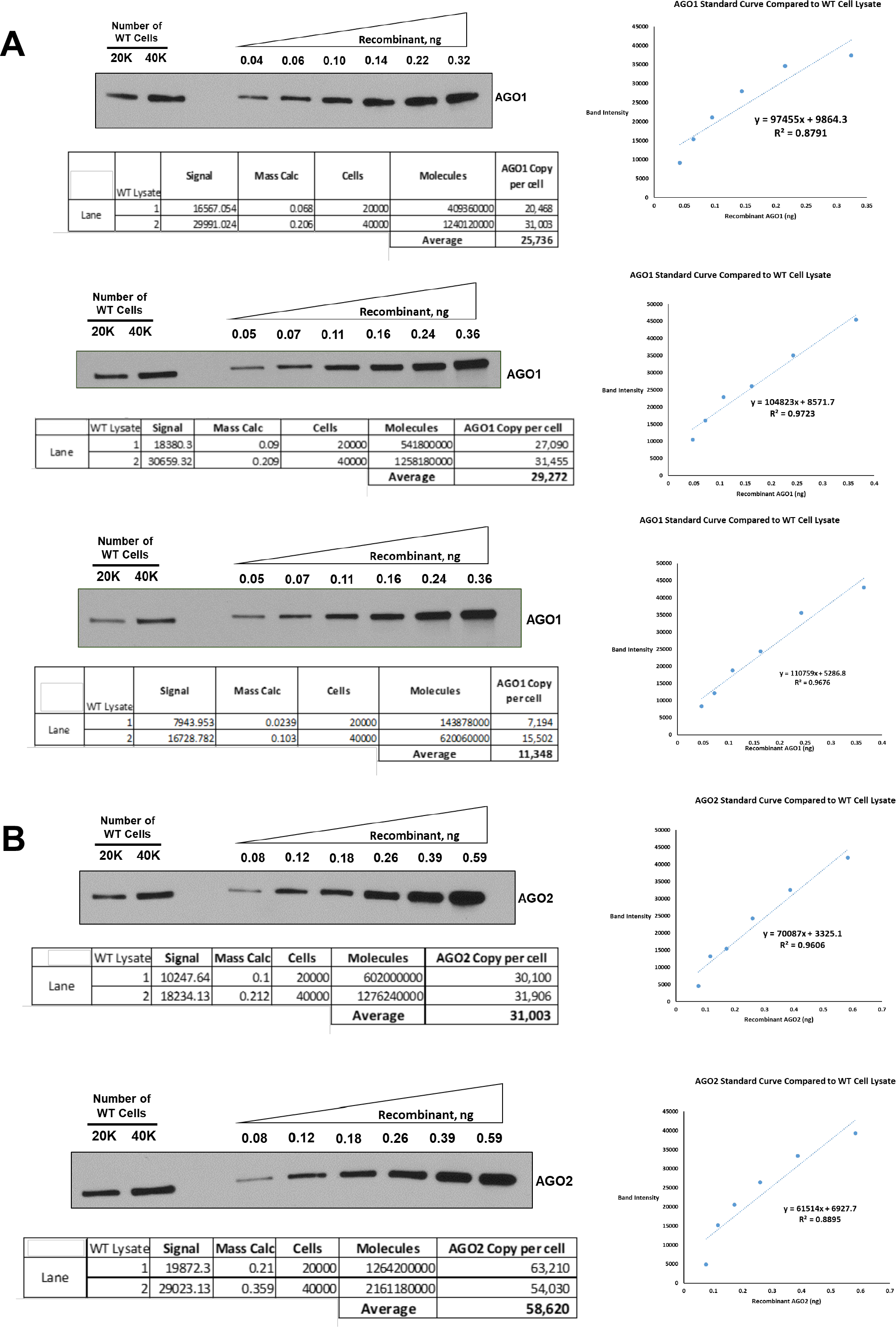
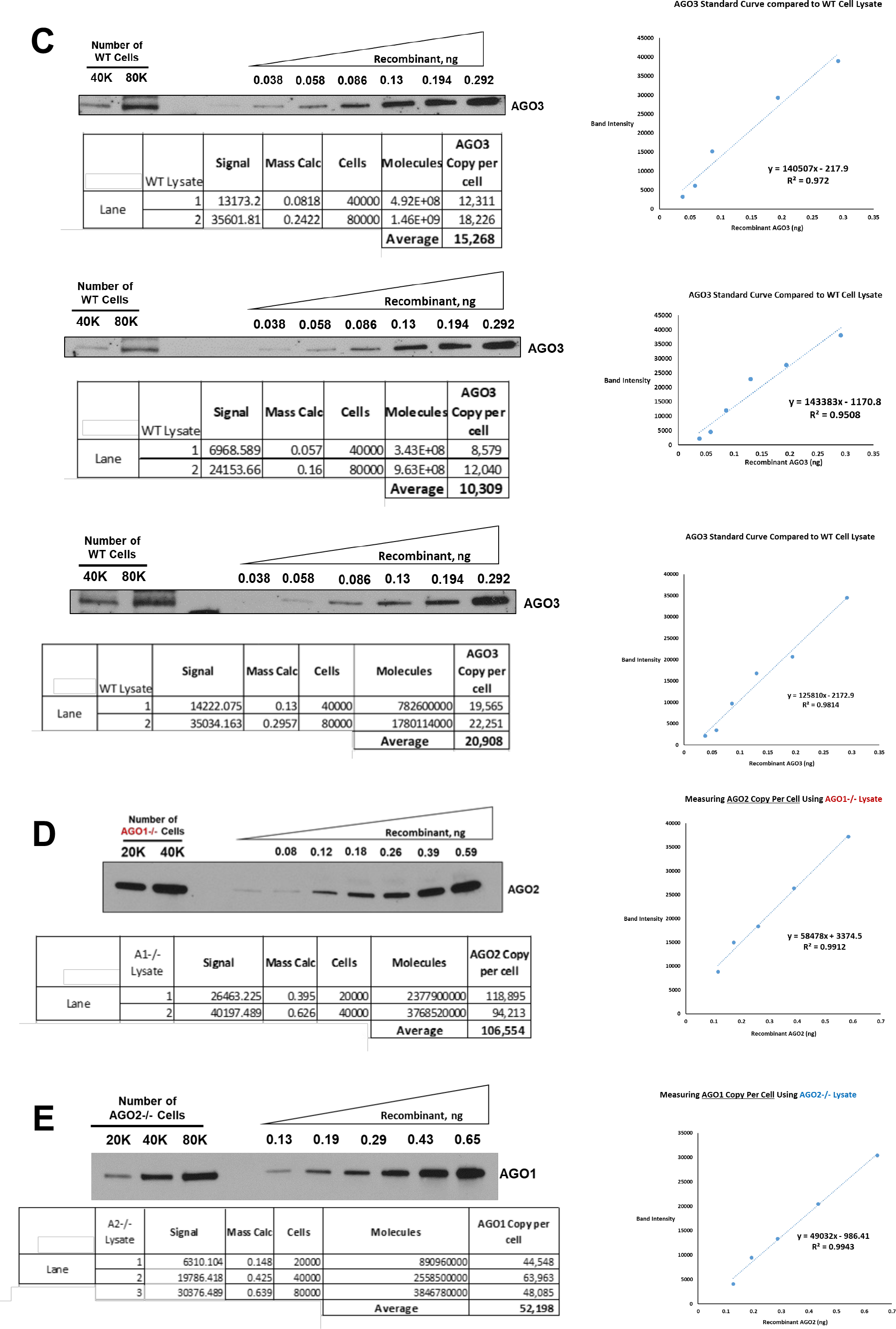
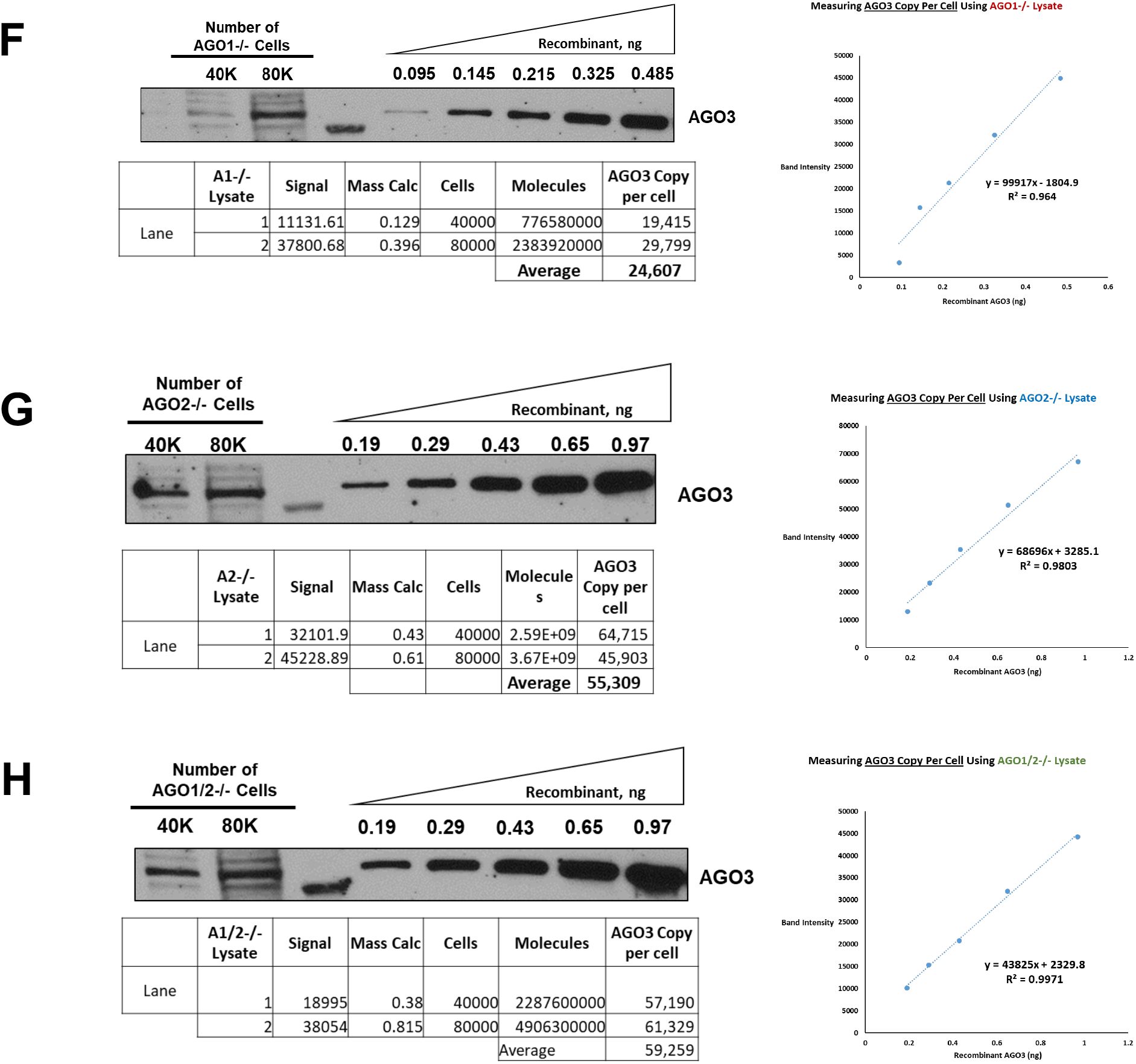
Absolute Quantification of Argonaute proteins in WT, AGO1−/−, AGO2−/−, AGO1/2−/−, and AGO1/23−/− HCT116 Cells. (A-H)Recombinant AGO1, AGO2, and AGO3 protein (Active Motif) were used to generate standard curves to estimate the mass of AGO1, AGO2, or AGO3 in a given quantity of HCT116 lysate corresponding to a known number of cells. Western blot signals were quantified using a digital scanner and ImageJ (NIH). Quantification summarized in tables below blots. (A) Quantification of AGO1 in WT cells. (B) Quantification of AGO2 in WT cells. (C) Quantification of AGO3 in WT cells. (D) Quantification of AGO2 in AGO1−/− cells. (E) Quantification of AGO1 in AGO2−/− cells. (F) Quantification of AGO3 in AGO1−/− cells. (G) Quantification of AGO3 in AGO2−/− cells. (H) Quantification of AGO3 in AGO1/2−/− cells.

**Supplemental Figure 4.**
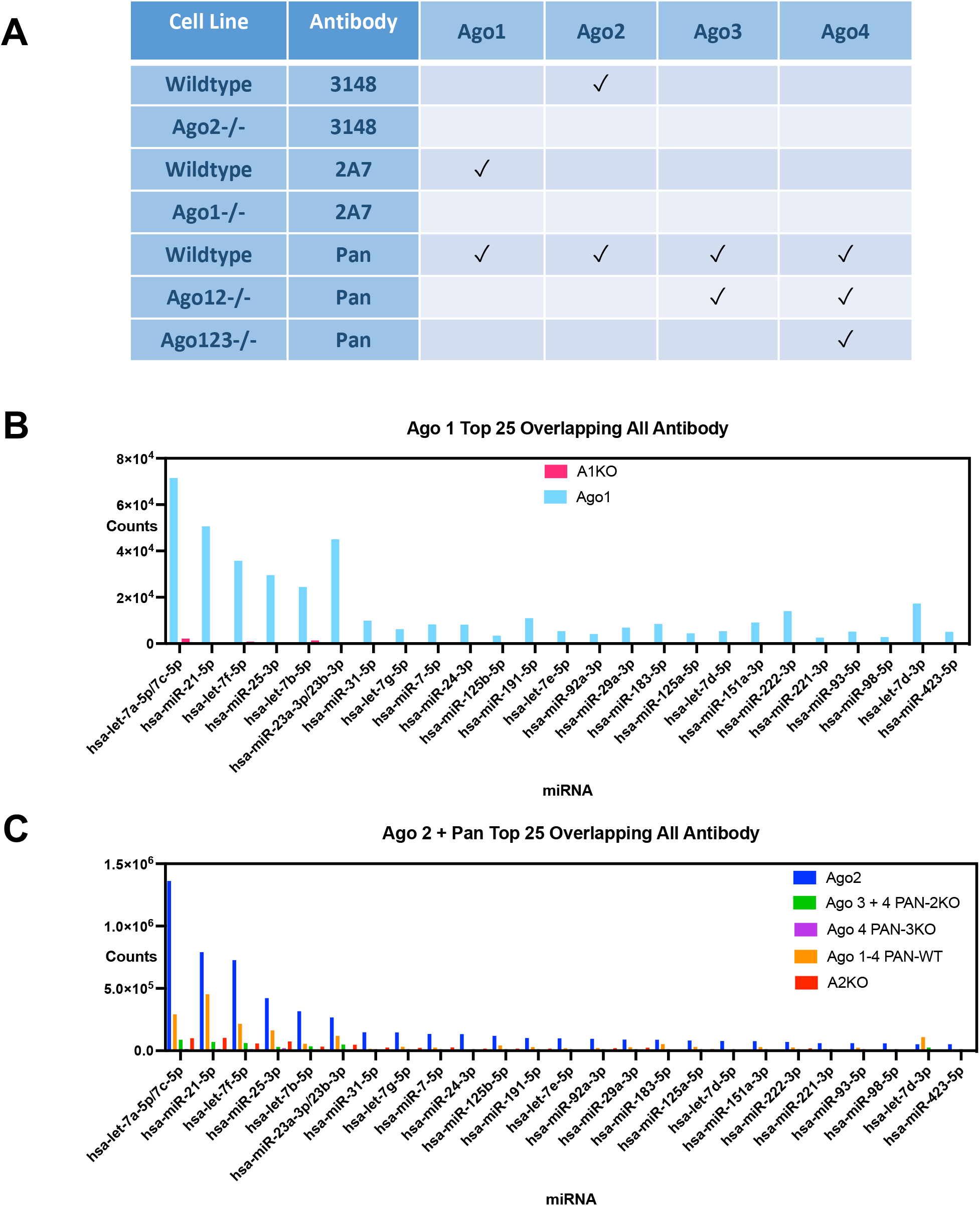
(A) Table describing the different experimental approaches to immunoprecipitate each Argonaute protein in WT and AGO−/− cells with different antibodies. (B) Chart of AGO1 counts for the top 25 of the 60 overlapping top 100 miRNA. (C) Chart of AGO2 and PAN antibody counts for the top 25 of the 60 overlapping top 100 miRNA.

**Supplemental Figure 5.**
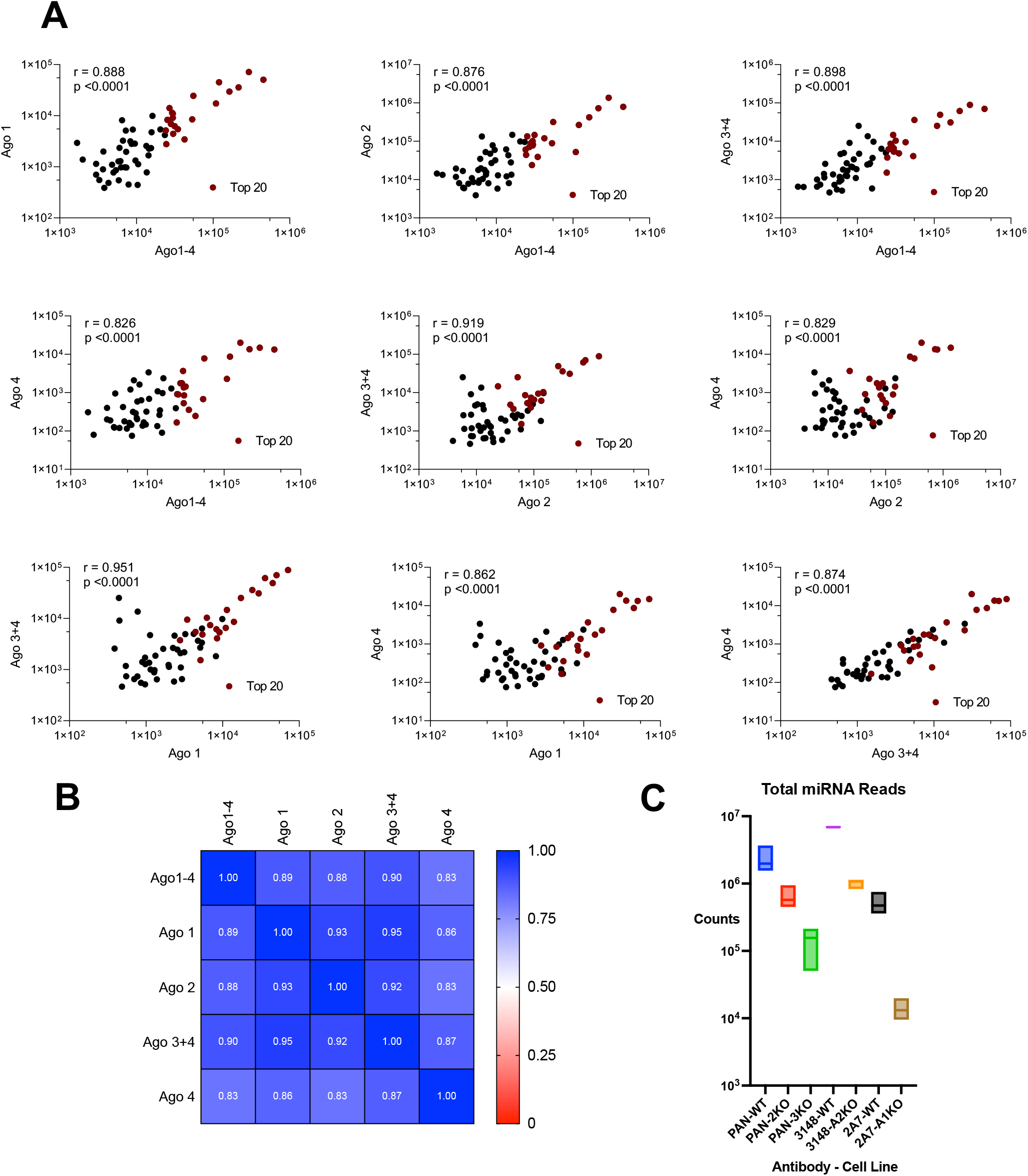
(A) Correlation plots of miRNA reads for the top 60 miRNAs found in all pulldowns across WT, AGO1−/−, AGO2−/−, AGO1/2−/− and AGO1/2/3−/− cell lines, where r is the Pearson correlation value and p is significance. The top 20 miRNAs with the highest read count from the anti-pan-AGO antibody pulldown are highlighted in red. (B) Heatmap of Pearson correlation values for all of the IP pulldowns. P<0.0001 for all correlations calculated. (C) Total mapped reads (counts) from each AGO immunoprecipitation.

